# Efficiency of Learned Indexes on Genome Spectra

**DOI:** 10.1101/2025.07.10.664199

**Authors:** Md. Hasin Abrar, Paul Medvedev, Giorgio Vinciguerra

## Abstract

Data structures on a multiset of genomic *k*-mers are at the heart of many bioinformatic tools. As genomic datasets grow in scale, the efficiency of these data structures increasingly depends on how well they leverage the inherent patterns in the data. One recent and effective approach is the use of learned indexes that approximate the rank function of a multiset using a piecewise linear function with very few segments. However, theoretical worst-case analysis struggles to predict the practical performance of these indexes.

We address this limitation by developing a novel measure of piecewise-linear approximability of the data, called CaPLa (Canonical Piecewise Linear approximability). CaPLa builds on the empirical observation that a power-law model often serves as a reasonable proxy for piecewise linear-approximability, while explicitly accounting for deviations from a true power-law fit. We prove basic properties of CaPLa and present an efficient algorithm to compute it. We then demonstrate that CaPLa can accurately predict space bounds for data structures on real data. Empirically, we analyze over 500 genomes through the lens of CaPLa, revealing that it varies widely across the tree of life and even within individual genomes. Finally, we study the robustness of CaPLa as a measure and the factors that make genomic *k*-mer multisets different from random ones.

**Supplementary Material** *Software (Source Code)*: https://github.com/medvedevgroup/CaPLaarchivedatswh:1:dir:da45f156bdafa582fd16f04690ee49e184bf3590

**Funding:** This material is based upon work supported by the NSF under Grants No. DBI2138585 and OAC1931531. Research reported in this publication was supported by the National Institute Of General Medical Sciences of the NIH under Award Number R01GM146462. The content is solely the responsibility of the authors and does not necessarily represent the official views of the NIH. GV was supported by the NextGenerationEU – National Recovery and Resilience Plan (Piano Nazionale di Ripresa e Resilienza, PNRR) – Project: “SoBigData.it - Strengthening the Italian RI for Social Mining and Big Data Analytics” – Prot. IR0000013 – Avviso n. 3264 del 28/12/2021

## 1 Introduction

Data structures on *k*-mers (fixed-length strings) originating from genomes form a crucial component of many bioinformatics tools [20]. Their space usage is now a major bottleneck at the forefront of biological discovery. For example, the Logan project needed two petabytes to index the 31-mers from the Sequence Read Archive [5]. Existing data structures often meet the space challenge by exploiting the non-uniform structure of genomic *k*-mer sets [4]. In doing so, they circumvent [1, 15, 21, 26] the theoretical worst-and average-case lower bounds, which offer a more pessimistic perspective [23].

A popular approach is based on learned indexes, a recent research direction that takes advantage of the internal patterns in the rank curve of a dataset [3, 8, 16, 30]. The rank of an element *x* in a multiset *S* is defined as the position of the first occurrence of *x* in the sorted list of *S*. A learned index is constructed from *S* and, given a query *k*-mer, returns an error-bounded prediction for its rank. Initial approaches used machine learning to construct the index [11, 12, 13], but it later turned out that in this setting, it is more effective to model the rank curve using a piecewise linear approximation [1, 2, 15], as depicted in Figure 1.

**Figure 1.**
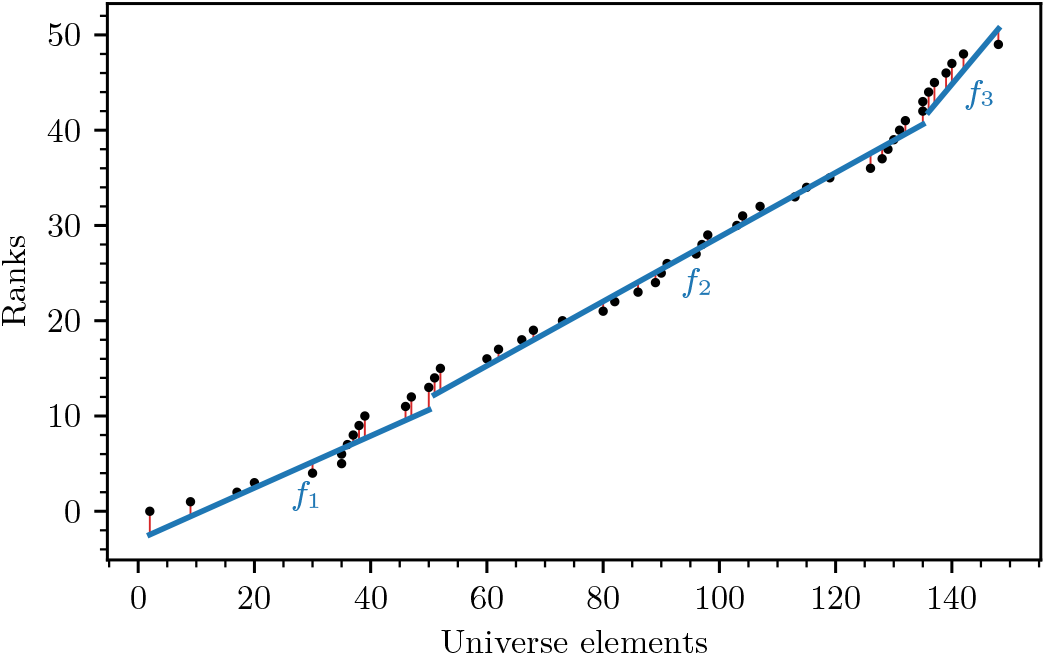
An example multiset *S* of size *N* = 50 with *n* = 48 distinct elements drawn from a universe of size *u* = 150, plotted as points { (*x*, rank_*S*_ (*x*)}_*x*∈*S*_ on the plane. The points are approximated using a PLA with error bound *ε* = 2, where the vertical orange lines show the error. The PLA uses 3 segments, which is the smallest possible for *S* at *ε* = 2, so *b*(*ε*) = 3.

Learned indexes based on piecewise linear approximations (PLAs) of the rank curve have resulted in practically groundbreaking time/space tradeoffs, both on genomic [1, 6, 15] and other kinds of data [6, 9, 10, 14, 17, 18, 19, 29]. However, existing theoretical worst-case analyses only show improved performance under assumptions that are unrealistic for genomic datasets, such as the independence and identical distribution of gaps between *k*-mers [7]. This leaves a gap between the theoretical analysis of *k*-mer-based learned indexes and their much better performance on real data [1, 15]. By better understanding and characterizing the structural properties of genomic *k*-mer sets, we can not only better predict the performance of learned indexes but also drive the development of new PLA-based tools that maximally exploit the properties of real data.

Our approach in this paper is to develop a measure of piecewise linear (PL) approximability which can be used to parameterize the theoretical analysis of PLA-based methods. Parameterizing string complexity is an active area of research [24], but none of the existing measures capture the PL-approximability of a genome’s rank curve. Previous papers have proposed modeling the PL-approximability of *S* using a power-law [7, 9], and have implicitly used the two parameters of a power-law fit as proxies for the PL-approximability of *S*. However, the interpretation of such parameters is limited when the rank curve strays from a power-law, as (we will show) occurs in real genomic data.

We develop a novel measure of PL-approximability, which we call CaPLa (Canonical PL-approximability) (Section 2). It builds on the idea of a power-law fit but adds the uncertainty of the fit as part of the measure. We prove basic properties of CaPLa, including its existence, and give an algorithm to compute it (Section 3). The algorithm is exact under certain conditions of the data, which we empirically show are nearly always met in practice. We demonstrate how CaPLa can be applied to derive space bounds that accurately reflect properties of real data (Section 4).

Finally, we apply CaPLa to analyze genome spectra, where a spectrum is the multiset of all *k*-mers appearing in a string.^1^ We use a dataset of more than 500 genomes and find that CaPLa varies greatly across the tree of life and even within individual genomes (Section 5). We also show that CaPLa is a good predictor of the space usage of a learned index data structure (the PLA-index [1]). We use controlled experiments to elucidate what makes genome spectra different from random *k*-mer multisets. Finally, we verify that CaPLa has some of the desired properties of a measure of genome complexity, namely robustness to variation in genome or *k*-mer size. Our work helps reduce the gap between theory and practice and aims to deepen our understanding of when and how to best apply PLA-based indexes to real genomic data.

## 2 PL-approximability definitions and motivation

In this section, we present the key definitions of PL-approximability and the motivation behind them. We start with a multiset *S* of elements from an integer universe [*u*] = {0, 1, …, *u*−1}.In the case of *k*-mers, *u* = 4^*k*^, though our definitions work for any *u*. We will use the notation that *n* is the number distinct and *N* is the total number of elements in *S*. The function we will be approximating is defined as RANK_*S*_(*x*) = |{ *y* ∈*S*| *y*≤}|, for all *x*∈[*u*]. The key tool in designing PLA-based learned data structures [1, 6, 9, 10, 15] is the following:

### ▸ Definition 1

(Piecewise linear *ε*-approximation)

*For a given positive integer ε, a* piecewise linear *ε*-approximation (PLA) *of S is a partition of* [*u*] *into subintervals such that, for each subinterval* [*a*_*i*_, *b*_*i*_], *there exists a segment (i*.*e*., *a linear function) f*_*i*_ *such that* |*f*_*i*_(*x*) − *rank*_*S*_(*x*)| ≤ *ε for all x* ∈ [*a*_*i*_, *b*_*i*_].

Figure 1 shows an example of a PLA. The usefulness of a PLA comes from the fact that a data structure can avoid storing the full rank function and instead just store the PLA, as long as it has a way to handle the uncertainty due to the error of the approximation. To this end, the PLA with the smallest number of segments is the most space-efficient representation. Such a PLA can also be computed efficiently [25]. As a result, minimum-sized PLAs have formed the basis of several data structures [1,6,9,17,19,29]. We formalize it with the following definition.

### ▸ Definition 2

(PLA-size) *The* PLA-size *of S is the function b* : Z^+^ → Z^+^ *mapping a positive integer ε to the smallest integer b*(*ε*) *such that there exists a piecewise linear ε-approximation of S using exactly b*(*ε*) *segments*.

In this paper, we introduce a closely related measure, but one that factors out the effect of the data size.

### ▸ Definition 3

(PL-approximability)*The* PL-approximability *of S is the function mapping a positive integer ε to* 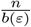, *where b*(*ε*) *is the PLA-size of S*.

The PL-approximability of *S* can be interpreted as the longest average number of elements spanned by a segment in a PLA of *S*, so it directly quantifies the efficiency of a PLA in capturing the structure and distribution of the underlying data.

The space and time performance of learned data structures based on PLAs are often bounded in terms of the PLA-size [1, 6, 9]. Typically, smaller values of *ε* lead to faster query times, but require a larger number of segments *b*(*ε*) and thus increased space. The PLA-size and PL-approximability can then be tabulated for a given dataset and plugged into these space-and time-bounds to assess the resulting space-time trade-off. However, this approach provides little insight into broader trends and differences among data distributions. Ideally, the PL-approximability could be modeled as a parametrized family of functions, such that fitting this family to *S* yields parameters that fully capture the PL-approximability of *S*.

A key difficulty is that the shape of the PLA-size function varies significantly depending on the structure and distribution of the underlying data. In general, the PLA-size is bounded between *b*(*ε*) = 1 and *b*(*ε*) = *N/*(2*ε*) for all *ε* [9]. *For sets where the gaps between elements are random and independently drawn, the PLA-size has been shown to be b*(*ε*) = 𝒪 (*N/ε*^2^) [7]. For real data, the PLA-size rarely admits a closed-form expression, but power-law relations of the form *b*(*ε*) = *n/*(*βε*^*α*^) have been empirically observed [7, 9]. Here, *β* and *α* are dataset-specific constants found through fitting to the tabulated PLA-size. This power-law modeling provides a degree of generality, but when the PLA-size does not exactly follow a power-law, it has several drawbacks. Figure 2 demonstrates that this type of modeling 1) has a high variability in the error, 2) is not robust to the choice of fitting algorithm, and 3) is unpredictable in whether it over-or under-estimates the true PLA-size. Such drawbacks make it challenging to incorporate this type of model into downstream analyses of data structures, especially since its predictions are neither worst-nor average-case bounds.

**Figure 2.**
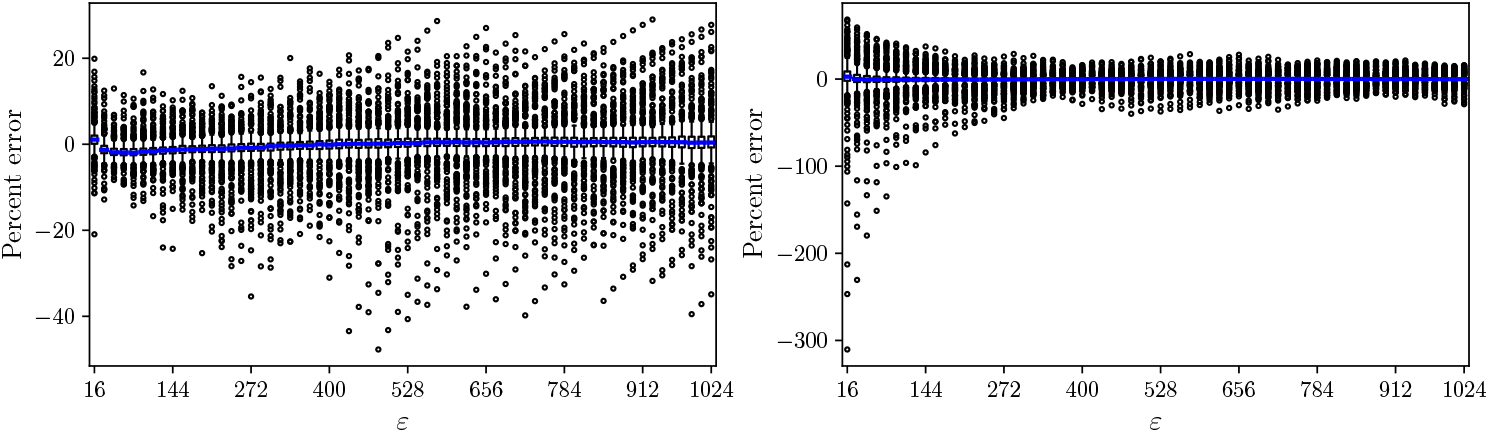
Errors due to fitting a single power-law curve to the PLA-size of a set of 513 genomes (dataset details are in Section 5.1). The left panel models the relationship as 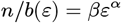 and uses Python’s *scipy*.*optimize*.*curve_fit* with the Levenberg–Marquardt algorithm to estimate *α* and *β*. The right panel uses the model log(*n/b*(*ε*)) = *α* log *ε*+log *β* and applies Python’s *scipy*.*stats*.*linregress* to estimate *α* and *β*. For each *ε* 16, 32, 48, 64, …, 1024, we calculate the percent error between *b*(*ε*) and the prediction *n/*(*βε*^*α*^) and show a box plot of the distribution aggregated over the 513 genomes (with the median line colored in blue).

To address this challenge, we propose bounding the PLA-size using two power laws instead of relying on a single fitted curve. We first fix a finite set *ε* ⊂ Z^+^ of *ε* values for which the PL-approximability is evaluated. This reflects practical applications, where *ε* is chosen from a set of values according to the desired space-time trade-off.

### ▸ Definition 4

(Bounded PL-approximability)*The PL-approximability of S is power-law bounded over ε with parameters* (*α, β*_low_, *β*_high_) *if*

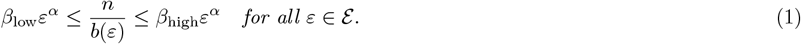

When *β*_low_ = *β*_high_, this is equivalent to a perfect power-law fit. The role of *α* is to capture the rate of segment growth as a function of *ε*. If *β*_low_ = *β*_high_, then when *ε* doubles, the average segment length increases by a factor of 2^*α*^. There is similarly an intuitive interpretation of *β*_low_ and *β*_high_: when *ε* = 1, the average segment length is between *β*_low_ and *β*_high_.

Definition 4 captures a full spectrum of data distributions and their PLA-size, from the best case *b*(*ε*) = 1 (with *α* = 0 and *β*_low_ = *β*_high_ = *n*), to the worst case *b*(*ε*) = *N/*(2*ε*) from [9] (with *α* = 1 and *β*_low_ = *β*_high_ = 2*n/N*), to the case of random gaps from [7] (with *α* = 2 and *β*_low_ = *β*_high_ = *cn/N* for a constant *c >* 0). More importantly, it enables bounding the PL-approximability of real datasets where an exact power-law fitting is not possible.

The PL-approximability of a single dataset can be power-law bounded for infinitely many values of *α, β*_low_, and *β*_high_. To capture a single parameterization that can be used as a proxy for the PL-approximability of a dataset, we introduce the following notion of *canonical PL-approximability*.

### ▸ Definition 5

(CaPLa) *A pinch point of S is any α*^∗^ *that satisfies*

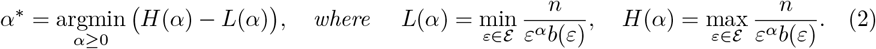

*If a unique pinch point α*^∗^ *exists, then we let* 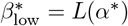 *and* 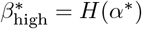 *and say that S has a canonical PL-approximability (CaPLa) of* 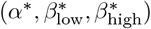. *Otherwise, we say that the CaPLa of S is undefined*.

We note that if *S* has CaPLa 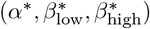, then Equation 1 immediately implies that *S* is power-law bounded with parameters 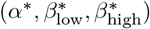

## 3 Existence and computation of the canonical PL-approximability

In this section, we show under which conditions the canonical PL-approximability exists and how it can be efficiently computed. We do so by introducing a geometric object, the *twisted ribbon*, which captures the entire family of valid power-law bounds for *S*. We then study the structural properties of this ribbon to identify a *pinch point*, which yields the canonical PL-approximability (Definition 5). Finally, we describe a two-phase algorithm that efficiently locates a pinch point based on the twisted ribbon’s geometry.

We first observe that, for any given *α*, the tightest possible power-law bound on the PL-approximability of *S* is given by choosing *β*_low_ and *β*_high_ as *L*(*α*) and *H*(*α*), respectively. The proof follows easily from the definitions of *L* and *H*.

▸ **Lemma 6***For any α* ≥ 0 *and δ>* 0, *we have that (i) S is power-law bounded over* E *with parameters* (*α, L*(*α*), *H*(*α*)), *and (ii) S is not power-law bounded over* E *with parameters* (*α, L*(*α*) + *δ, H*(*α*) − *δ*).

We can plot the functions *H* and *L* on a Cartesian plane with *α* along the horizontal axis and *β* along the vertical axis. The region enclosed between these two curves forms the aforementioned twisted ribbon, depicted in Figure 3 and formally defined as follows.

**Figure 3.**
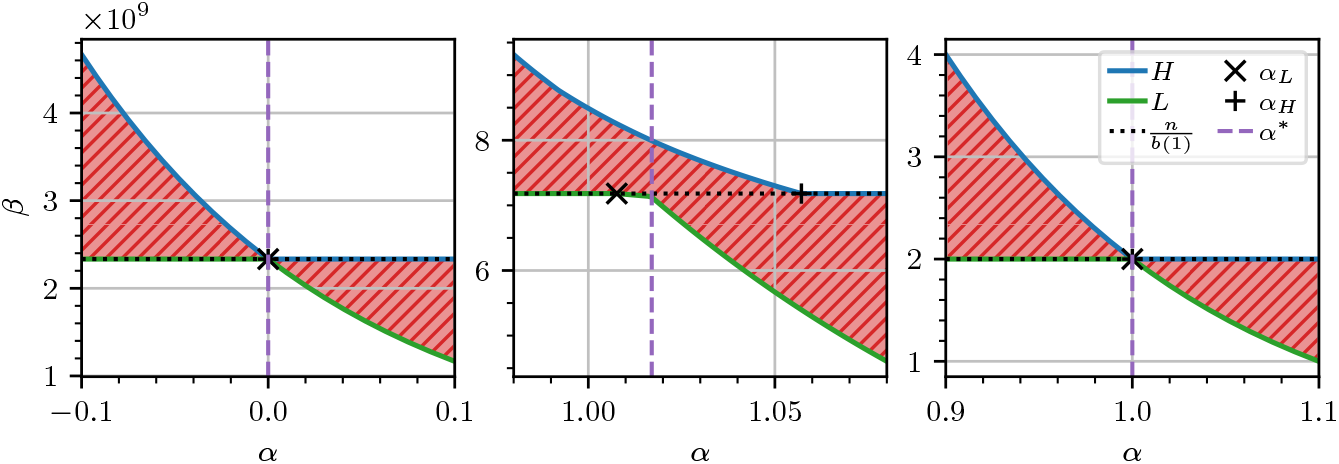
On the left, a twisted ribbon corresponding to a set *S* giving the best case *b*(*ε*) = 1 for any *ε* ∈{1, …, 1024}. On the center, a twisted ribbon derived from the human genome (T2TCHM13v2.0). On the right, a twisted ribbon corresponding to a set *S* giving the worst case *b*(*ε*) = *N/*(2*ε*) for any *ε*. For symmetry with the other plots, the left plot extends to values of *α <* 0 despite the ribbon being formally defined for *α* ≥ 0.

### ▸ Definition 7

(Twisted ribbon)*The twisted ribbon of S is the region in the α-β space enclosed between the curves L*(*α*) *and H*(*α*) *for any α >* 0. *Formally, the twisted ribbon of S is the set* 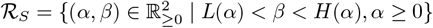.

Observe that choosing *β*_low_ or *β*_high_ inside of the twisted ribbon (that is, setting *β*_low_ *> L*(*α*) or *β*_high_ *< H*(*α*) for some *α*) would violate Equation 1, and therefore would not yield a valid bound on the PL-approximability of *S*. This reduces the space of relevant power-law bounds to that of {(*α, L*(*α*), *H*(*α*))} _*α*≥ 0_.

We further aim to summarize this entire family of bounds into a single representative instance. Intuitively, this means identifying a value of *α* that yields the tightest bound. This corresponds to a value *α*^∗^ that minimizes the twisted ribbon width *W* (*α*) = *H*(*α*)−*L*(*α*), i.e., what we call a pinch point (Definition 5).

To study the existence and computation of a pinch point, we first show some structural properties of the twisted ribbon. For technical convenience, we assume that the (nonincreasing integer) function *b*(*ε*) has been extended to a continuous, nonincreasing, and differentiable function on the reals, e.g., via monotone cubic interpolation.

### ▸ Lemma 8

*If* 1 ∈ ε *and* |*ε*| *>* 1, *then:*

1. *H and L are continuous functions*.
2. *For any α* ≥ 0, *it* 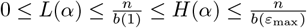, *where ε*_max_ = max*ε*.
3. *There exists a value α*_*L*_ *such that* 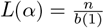, *and L*(*α*_*L*_ + *δ*) *decreases as δ>* 0 *increases. We call α*_*L*_ *a flattening point of S*.
4. *There exists a value α*_*H*_ *such that* 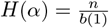, *and H*(*α*_*H*_ − *δ*) *increases as δ>* 0 *increases. We also call α*_*H*_ *a flattening point of S*.
5. *The flattening points satisfy α*_*L*_≤*α*_*H*_.

Note that the lemma holds for any dataset, as its conditions are related to a parameter (ε) chosen by the user. The following result shows that a pinch point exists and is confined between (or at) the flattening points mentioned in Lemma 8 and depicted in Figure 3.

▸ **Theorem 9** *If* 1 ∈ ε |ε| *and >* 1, *then a pinch point α*^∗^ *of S exists and is guaranteed to lie in the interval* [*α*_*L*_, *α*_*H*_], *where α*_*L*_ *and α*_*H*_ *are the flattening points of S*.

We now focus on finding a pinch point, that is, on solving the problem argmin _*α*_≥_0_ *W* (*α*). Observe *W* is non-differentiable due to the presence of min and max operations. Moreover, we assume that *W* is *unimodal*, namely, it has a unique minimum *α*^∗^, is strictly decreasing for all *α < α*^∗^, and strictly increasing for *α > α*^∗^. While we do not have any general theoretical guarantee on the unimodality of *W* (except in simple cases such as those shown in the left and right plots of Figure 3), our empirical observation suggests that unimodality often holds in practice (see Section 5.5). If this is not the case, our algorithm finds a local minimum instead. We also assume *b*(*ε*) has been precomputed for all *ε* ∈ ***ε***, allowing *W* (*α*) to be evaluated in 𝒪 (|ε|) time via a direct computation based on Equation 2. This precomputation can be done in 𝒪 (*N*|ε|) time [25].

The algorithm works in two phases. In the first phase, we identify the flattening points *α*_*L*_ and *α*_*H*_ by observing from Lemma 8 that *α*_*L*_ is the smallest value of *α* such that 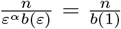, for some *ε* ∈ ε \ {1}. This equality simplifies to 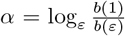, so we obtain 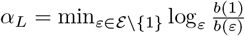, which can be computed in 𝒪 (|ε |) time. A similar derivation with max instead of min yields *α*_*H*_.

In the second phase, we run a derivative-free minimization algorithm (golden-section search) within the interval [*α*_*L*_, *α*_*H*_], which is guaranteed to contain *α*^∗^ due to Theorem 9. It converges in 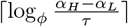 iterations, where *τ* is the desired tolerance, and *ϕ*≈1.618 is the golden ratio [27]. Since the cost of each iteration is dominated by the evaluation of *W* (*α*), the overall time complexity of finding the pinch point is 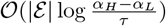. This gives the following theorem.

### ▸ Theorem 10

*Suppose the ribbon width W* (*α*) = *H*(*α*) − *L*(*α*) *is unimodal*, 1 ∈ ε, *and* |ε|*>* 1. *Suppose the value b*(*ε*) *has been precomputed for all ε* ∈ ε *in* 𝒪 (*N* |ε|) *time. Then, there exists an algorithm that finds the pinch point α*^∗^ *in* 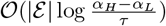 *time, where τ is the desired tolerance*.

Section 5.1 will show that the runtime of this algorithm is on the order of a few milliseconds in practice.

## 4 Using PL-approximability bounds in downstream applications

There are a number of PLA-based learned data structures whose space-time performance is bounded in terms of the PLA-size, such as the PGM-index [9, Theorem 1], LeMonHash [6, Theorem 2], and the PLA-index [1, Theorem 1]. In this section, we showcase how our PL-approximability framework can be used to reinterpret such bounds, focusing on the PLA-index as a concrete example particularly relevant in genomic applications. The PLA-index has been applied to substantially improve the run time of suffix array search and decrease the space of a read aligner and direct-access lookup tables [1].

The PLA-index uses a piecewise linear *ε*-approximation to a multiset of *k*-mers *S* to estimate the rank of a query *k*-mer in *S* [1]. *The tuning parameter ε* represents a time/space trade-off, with smaller *ε* values resulting in faster index query times but more segments.

Theorem 1 in [1] describes the size of the PLA-index as a function of the number of segments used by the PLA index (denoted by *s*_*ε*_), *N*, and *ε*:

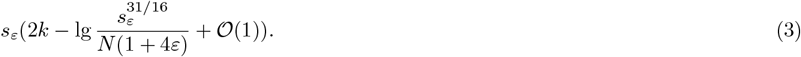

As *s*_*ε*_ can be high in the worst case, Equation 3 does not capture the fact that the space is very low in practice.

Here, we show that if *S* has a power-law bounded PL-approximability, then Equation 3 can be turned into a useful predictor of PLA-index space usage. One of the technical difficulties is that PL-approximability gives bounds on the PLA size *b*(*ε*), whereas Equation 3 is stated in terms of *s*_*ε*_. Though the PLA-index uses a segmentation based on the optimal one, it requires consecutive segments to share an endpoint and in some rare cases breaks segments. This dependence of *s*_*ε*_ on *b*(*ε*) is difficult to quantify in terms of *S*. Instead, we isolate the dependence by expressing our results in terms of 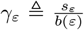. Though *γ*_*ε*_ is theoretically unknown (except that *γ*_*ε*_≥1), empirical results show that it is close to one on genome spectra. For example, for the 21-spectrum of chromosome 1 of the human genome, *γ*_*ε*_ values are below 1.21 for all tested *ε* up to *ε* = 1023.

▸ **Theorem 11***Let S be a multiset of k-mers, with n distinct elements, N total elements. Suppose the PL-approximability of S is* (*α, β*_low_, *β*_high_) *power-law bounded over* ε. *Let B*_*ε*_ *be the number of bits used by the PLA-index on S with tuning parameter ε. Then, for all ε* ∈ ε, *we have that*

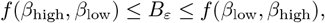

*Where*

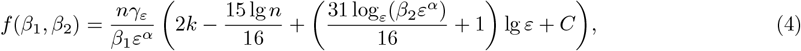

*and* 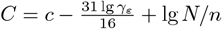 *and, for large enough* 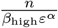, *c is a number between 5 and 7*.

We have isolated and combined some terms into a variable *C*, in order to convey that they can be thought of as constants in practice. First, *N/n* is in practice smaller than 2 when avoiding small values of *k*, for which a suffix array would not make sense as an application anyway. Second, as we mentioned earlier, *γ*_*ε*_ is in practice a number close to 1.

Interpreting Eq. 4 is helped by the observation that given the type of values we see in practice, the dominating term is the leading one, i.e. 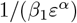. This intuition is confirmed by Figure 4AB, which plots Eq. 4 using real numbers from human chromosome 1. As suggested by the leading term, increasing the *α* value of a dataset by *δ*results in the leading term decreasing by a factor of *ε*^*δ*^, with the plots confirming this type of exponential decay for Eq. 4 overall. Note that for *ε* = 63 (Panel A), the decay is slower than for *ε* = 1023 (Panel B), as expected from the leading term. Figure 4CD shows how the gap between *β*_low_ and *β*_high_ affects the gap between the upper and lower bounds of Theorem 11 and gives the scale of the gap at which our bounds become meaningless.

**Figure 4.**
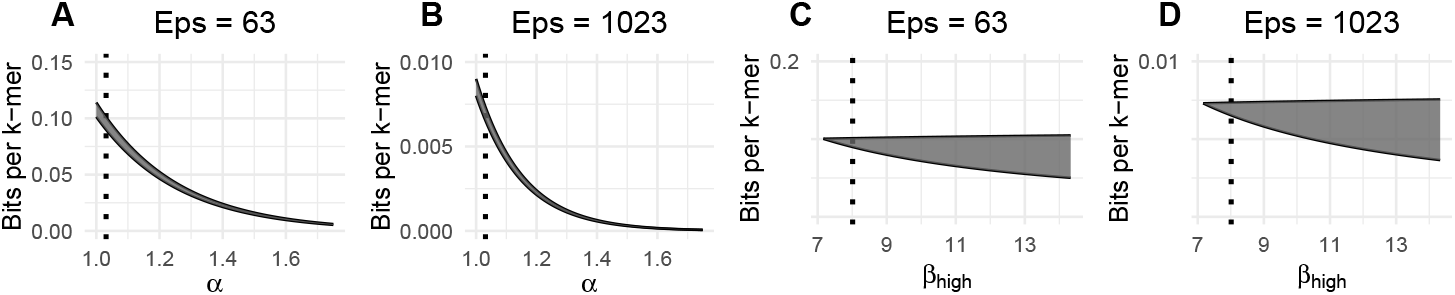
Analysis of space usage of PLA index, as predicted by Theorem 11, plugging in *C* = 6. In each panel, the shaded area is bounded from above by the upper bound curve and below by the lower bound curve. We use values corresponding to human chromosome 1 with *k* = 21: *n* = 195, 735, 278, *α*^∗^ = 1.03, 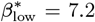, and 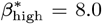, *γ*_63_ = 1.13 and *γ*_1023_ = 1.15. In panels A and B, we fix 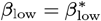 and 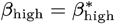 while varying *α*. The vertical dotted line corresponds to *α* = *α*^∗^. In panels C and D, we fix the values of *α* = *α*^∗^ and 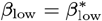 and vary *β*_high_. The vertical dotted line corresponds to 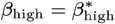

## 5 Experimental results

The goal of our experiments is threefold. First, we want to understand how much PL-approximability varies among the tree of life. Second, we want to understand the properties of genome spectra that make them different from random *k*-mer multisets. Finally, we want to evaluate the suitability of CaPLa as a measure of PL-approximability. Our tool for computing CaPLa, as well as reproducibility information, is available through GitHub at https://github.com/medvedevgroup/CaPLa.

### 5.1 Experimental setup

#### Data

We downloaded a sample of RefSeq genomes that are complete and full, do not have any missing bases, and are longer than 10,000nt. We further filtered out any genomes for which the number of segments at *ε* = 1024 was 1 (i.e. *b*(1024) = 1). This excludes genomes that are too trivial to offer meaningful insight and for which the PL-approximability becomes trivially dominated by *n*. We process the genomes to make everything uppercase, remove any letters that are not A, C, G, or T, and concatenate all contigs into a single string. The resulting dataset contains 513 genomes, representing the kingdoms of virus (91 genomes), bacteria (200 genomes), archaea (200 genomes), and fungi (22 genomes). The median genome length was 2.98 mil and the maximum was 63 mil, with the length distribution shown in further detail in Figure A.1. We also downloaded six other larger genomes: *C. elegans, P. yoelii, M. commoda, O. lucimarinus*, and the latest high-quality assemblies of the human (T2TCHM13v2.0) and Gorilla genomes (length information shown in Table A.1).

#### Tool

Our tool to compute CaPLa takes a genome, a *k*-mer size, and ε, builds a suffix array of the genome using libdivsufsort [22], computes the values {*b*(*ε*)}_*ε*∈ε_ using O’Rourke’s algorithm [25] on the genome’s *k*-mers (extracted in sorted order using the suffix array), and then computes CaPLa using the algorithm of Theorem 10 with tolerance *τ* = 1.49 × 10^−8^ (i.e., the square root of the machine epsilon). We use ε as the set of all integers from 1 to 1024, and *k* = 21 by default. We also construct the PLA-index [1] for *ε* ∈ {32, 64} for all genomes, using *k* = 21 as in previous works [1, 15]. Full details, including the accession number of the genomes, their CaPLa values, PLA-index sizes, and other associated information are available at our GitHub repository.

#### Runtime

For runtime analysis, we used a machine with a 2.10 GHz Intel Xeon CPU E5-2683 v4 processor with 512 GB of memory. To give a sense of the execution time of our pipeline, for a RefSeq genome of median length (GCF_001870125, 2.98M bases), it takes 0.318 s to construct the suffix array, 730.979 s to compute the values {*b*(*ε*)}_*ε*∈ε_, and seconds to run the algorithm of Theorem 10. Thus, the runtime of the algorithm of Theorem 10 is negligible and we do not investigate it further. A more detailed analysis of the runtime of computing *b*(*ε*) values is covered in other papers (e.g. [1]); since it is not a contribution of our work, we do not expand on those studies here.

#### CaPLa comparison

At first glance, comparing CaPLa values across genomes may seem challenging, as *α*^∗^ has a larger impact on the PL-approximability than 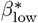 or 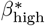, due to its presence in the exponent of *ε* in Equation 1. However, the values are easier to compare when the user has a desired query time, i.e. when we think of *ε* as being fixed. In this case, we define the *ε-mapped CaPLa* as the following two values: *ρ*_low_ = *α* + log_*ε*_ *β*_low_ and *ρ*_high_ = *α* + log_*ε*_ *β*_high_. The PL-approximability power-law bound (Equation 1) can then be rewritten in terms of *ρ*_low_ and *ρ*_high_, i.e. *ε*^*ρ* low;^ ≤ *n/b*(*ε*) ≤ *ε*^*ρ* high^. This replaces the three variables of CaPLa with two variables which, moreover, are now directly comparable between genomes. We can further average *ρ*_low_ and *ρ*_high_ to get the *averaged ε-mapped CaPLa*, which becomes useful for looking at CaPLa distributions across large datasets.

### 5.2 Understanding the scale of CaPLa

Table 1 shows the CaPLa values for our data. But, to make sense of the variability, we need to first understand how variations in CaPLa values translate into variations in the space of downstream data structures. The theoretical predictions of Theorem 11 allow such an analysis for the case of the PLA-index. To simplify the analysis, we consider the case that PL-approximability follows a perfect power law, i.e. there is a *β*^∗^ such that 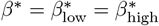. As Theorem 11 and Figure 4 indicated, the total bits per *k*-mer is approximately 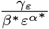 When *α*^∗^ increases by an additive factor of *δ*, then the approximate space decreases by a multiplicative factor of 1−*ε*^−*δ*^; when *β*^∗^ increases by a multiplicative factor of *δ*, then the approximate space decreases by a multiplicative factor of 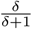. Table 2 shows these numbers for some concrete values that we encounter in our data. For example, in the bacterial genomes, the *α*^∗^ values increase by 0.1 from the 5th and 95th percentiles (Table 1); Table 2 shows that this corresponds to a decrease in space of 32% for the PLA index with *ε* = 64. Thus the variability in CaPLa across RefSeq corresponds to a substantial variability in predicted bits per *k*-mer of the associated PLA-index.

**Table 1.** CaPLa of the RefSeq dataset and of six additional genomes. The first row corresponds to all of the RefSeq dataset, while the later four rows separate the dataset into Kingdoms. Note that percentiles are not reported for the last six rows because each row corresponds to a single genome.

| Category | $\alpha^*$ | | | $\beta_{\text{low}}^*$ | | | $\beta_{\text{high}}^*$ | | |
| --- | --- | --- | --- | --- | --- | --- | --- | --- | --- |
|  | 5th% | Mean | 95th% | 5th% | Mean | 95th% | 5th% | Mean | 95th% |
| RefSeq dataset | 1.03 | 1.09 | 1.16 | 4.8 | 7.4 | 8.6 | 7.1 | 9.0 | 11.1 |
| Virus | 1.06 | 1.12 | 1.19 | 3.4 | 5.6 | 7.9 | 7.1 | 8.6 | 10.8 |
| Bacteria | 1.03 | 1.08 | 1.12 | 6.4 | 7.8 | 8.6 | 7.0 | 9.1 | 11.2 |
| Archaea | 1.03 | 1.07 | 1.12 | 6.7 | 7.8 | 8.6 | 7.2 | 8.9 | 10.7 |
| Fungi | 1.08 | 1.12 | 1.17 | 6.5 | 8.2 | 9.0 | 7.3 | 10.3 | 12.5 |
| *C. elegans* |  | 1.06 |  |  | 7.4 |  |  | 8.2 |  |
| *P. yoelii* |  | 1.00 |  |  | 5.5 |  |  | 6.1 |  |
| *M. commoda* |  | 1.06 |  |  | 7.4 |  |  | 8.2 |  |
| *O. lucimarinus* |  | 1.06 |  |  | 7.4 |  |  | 8.1 |  |
| Human |  | 1.02 |  |  | 7.1 |  |  | 8.0 |  |
| Gorilla |  | 1.00 |  |  | 7.1 |  |  | 7.9 |  |

**Table 2.** The effect of CaPLa on PLA-index space. We show the decrease in bits per distinct *k*-mer when adding *δ*to *α* (left columns) and multiplying *β* by (1 + *δ*) (right columns). The numbers without parenthesis indicate the predicted decrease using 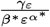, the leading term of Equation 4. The numbers in parenthesis indicate the change predicted by the full form of Theorem 11, using *C* = 6 and the real values from human chromosome 1 with *k* = 21 (*n* = 195, 735, 278, *α*^∗^ = 1.0309, 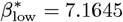, and 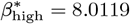, *γ*_63_ = 1.13 and *γ*_32_ = 1.13).

| | $\alpha \leftarrow \alpha + \delta$ | | | $\beta \leftarrow \beta(1 + \delta)$ | |
| --- | --- | --- | --- | --- | --- |
| | $\delta = 0.01$ | $\delta = 0.05$ | $\delta = 0.1$ | $\delta = 0.1$ | $\delta = 0.5$ |
| $\epsilon = 32$ | 3% (3%) | 16% (15%) | 29% (28%) | 9% (9%) | 33% (32%) |
| $\epsilon = 64$ | 4% (4%) | 19% (18%) | 34% (32%) | 9% (9%) | 33% (32%) |
| $\epsilon = 1024$ | 7% (6%) | 29% (28%) | 50% (48%) | 9% (9%) | 33% (32%) |

### 5.3 Variation of CaPLa among the tree of life

To better understand the distributions of CaPLa among different kingdoms, Figure 5AC shows the distributions of the averaged *ε*-mapped CaPLa. The fungi kingdom has a noticeably higher CaPLa. The difference is substantial, as is reflected in the lower PLA-index space usage of fungi genomes compared to the other kingdoms (Figure 5BD). The reasons for this could be due to biological factors but could also be due to biases in the type of genomes that have been sequenced and/or deposited into RefSeq.

**Figure 5.**
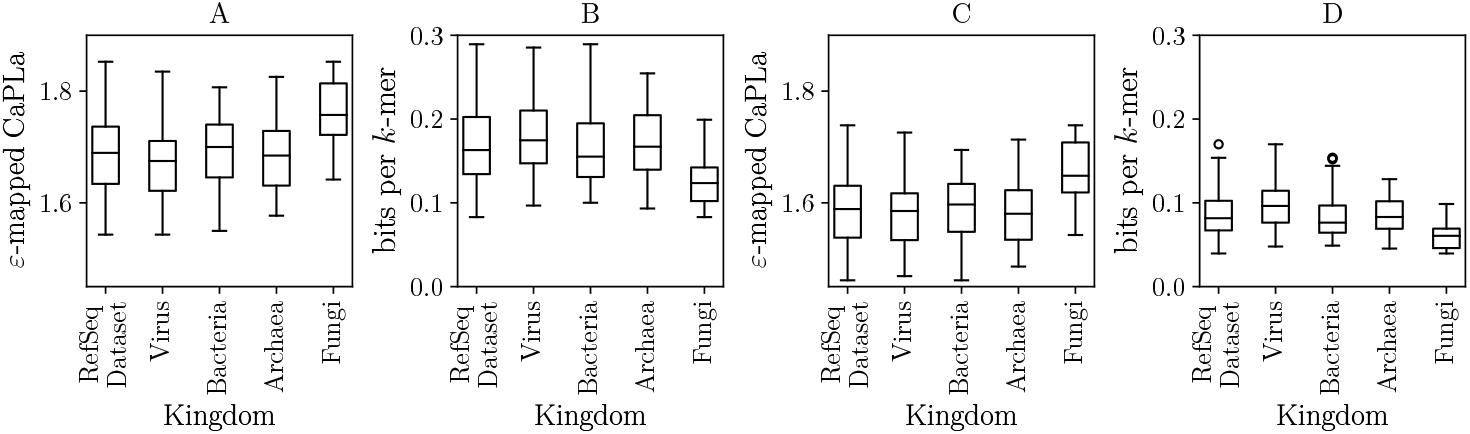
Distribution of averaged *ε*-mapped CaPLa across RefSeq. Panels A and C show the values for *ε* = 32 and *ε* = 64, respectively. Panels B and D show the distribution of bits per distinct *k*-mer in the constructed PLA indexes. In each panel, the left box plot is for our whole RefSeq dataset, while the remaining four box plots correspond to subsets according to Kingdoms.

Figure 6 shows the space usage of the PLA index and breaks down the relationship between bits per distinct *k*-mer and the CaPLa separately for each genome. The variation is substantial, with space differences as big as threefold between different RefSeq genomes. Furthermore, the figure shows a strong correlation between CaPLa and bits per distinct *k*-mer of the PLA-index.

**Figure 6.**
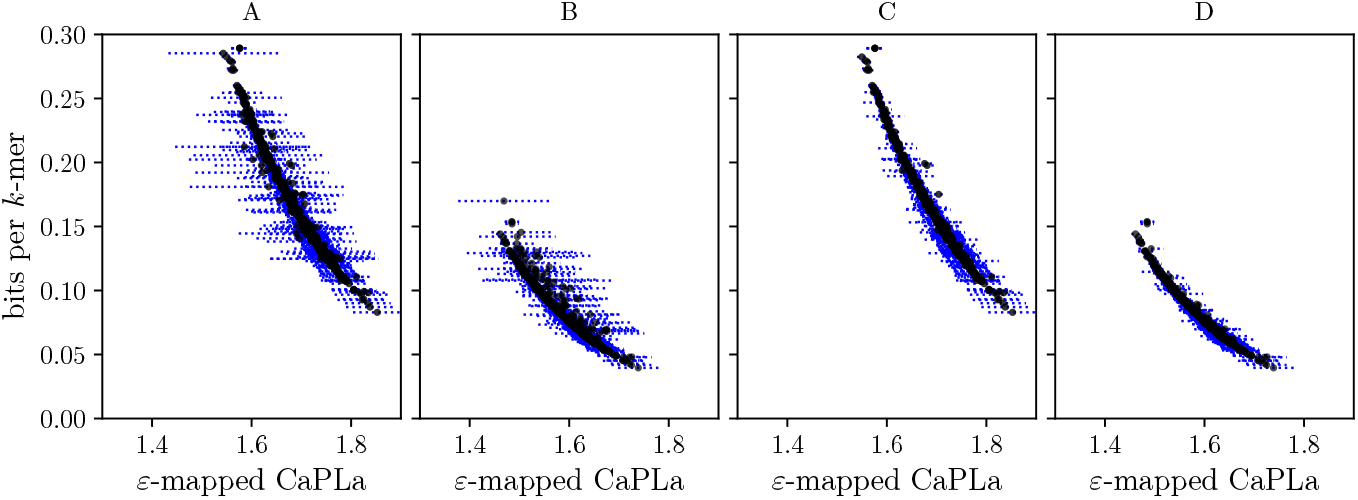
Relation between the bits per distinct *k*-mer of the PLA-index to the *ε*-mapped CaPLa for the RefSeq dataset. Panel A shows the result for *ε* = 32 and Panel B for *ε* = 64. Each dashed horizontal line represents the range of the *ε*-mapped CaPLa for a single genome and each black dot represents an averaged *ε*-mapped CaPLa for a single genome. Panels C and D show the same information as panels A and B, but we additionally filter out all genomes that are shorter than 80, 000nt. For such shorter genomes, the constant overhead of the PLA-index data structures starts to dominate the part that makes use of the PL-approximation; as the plot shows, most of the widest horizontal lines in panels A and B are gone after filtering out the shorter genomes.

We also split the human genome into its 24 constituent chromosome sequences. Figure 7 shows that CaPLa varies substantially across the chromosomes, indicating that PL-approximability varies not only between different species but also within the genome of a single species. Figure 7 also shows a similar pattern for the chromosomes of the Gorilla genome. Chromosome Y, which is known for being highly repetitive, is an outlier in both species, with substantially lower CaPLa than other chromosomes.

**Figure 7.**
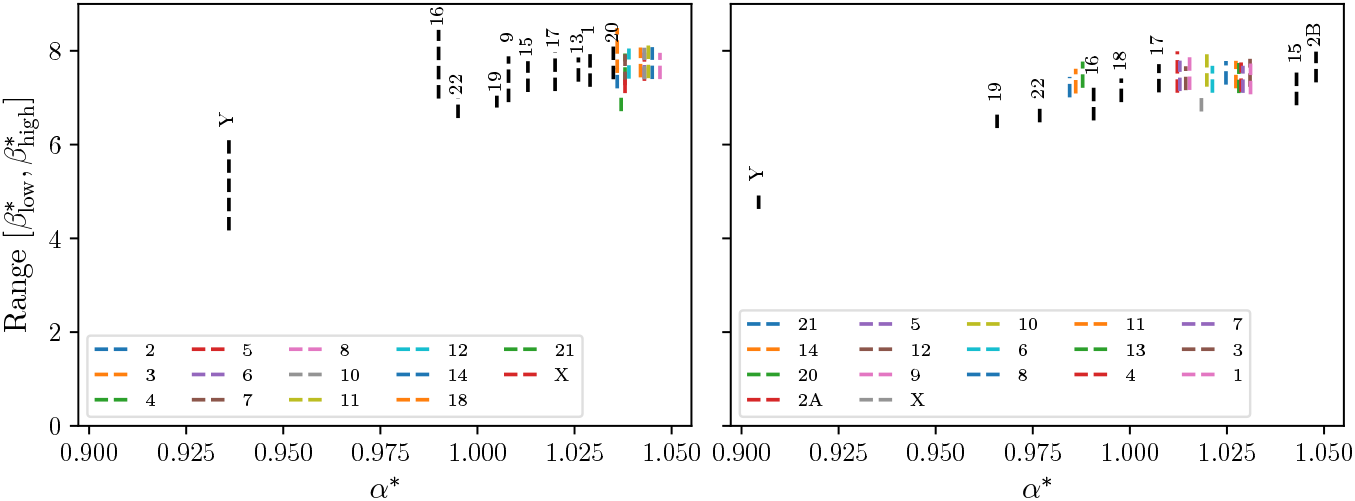
CaPLa of Human (left) and Gorilla (right) chromosomes. Each vertical segment is labeled with the corresponding chromosome, either in the legend or directly above the segment.

In spite of the CaPLa variability, our results give a general range of CaPLa for genomes that are not part of our dataset. This gives a user with a new genome sequence a robust estimate of the expected efficiency of a PLA-based learned index. This is in contrast with datasets from other domains that have been shown to roughly fit a power-law curve with significantly different *α* values, such as *α* = 1.8 on book popularity data, *α* = 1.4 on Facebook user IDs, and *α* = 2 on random gaps [7].

### 5.4 What makes a genome spectrum special?

Figure 8 shows a huge difference between the CaPLa of human chromosome 1 (*α*^∗^ = 1.03) and a random multiset of *k*-mers of the same size (*α*^∗^ = 1.86). Theorem 11 predicts a 26-fold difference in the size of the corresponding PLA indexes, at *ε* = 64. We therefore wanted to tease out the effect of several factors which distinguish a genome spectrum from a random multiset. The first factor is that the *k*-mers of a genome spectrum must be overlapping, i.e. not all multisets of *k*-mers can be put together into a continuous genome sequence, even allowing for chromosome breaks. To test the effect of this, we create a random string with the same length as chr1 and find that it also had *α*^∗^ = 1.86. Thus, this factor does not seem to play a role, at least in isolation.

**Figure 8.**
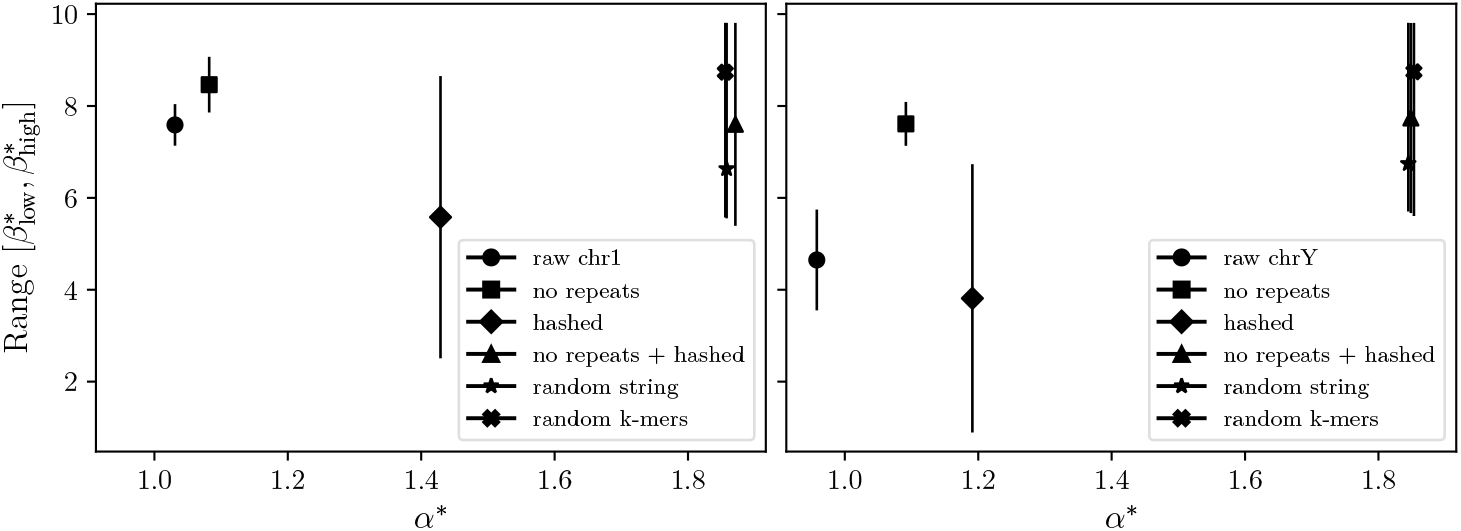
PL-approximability of a genome spectrum versus a random *k*-mer multisets. The left and right panels show human chromosomes 1 and Y, respectively, for *k* = 21. The y-axis shows the range from 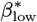 to 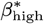 The random string and the random *k*-mers are generated so as to match the number of *k*-mers in the raw sequence.

A second effect is that chr1 is not random but is evolved to have a biological function. One consequence is that the genome has *k*-mers which appear more than once, which is rare in a random string of the same length (for *k* = 21). To test the effect of this, we make a linear scan of chr1 and clip out any encountered *k*-mer which we have previously seen. In this way, we create a genome-like string which is nearly repeat-free. We found *α*^∗^ = 1.08 for this low-repeat string, suggesting that repeats do negatively affect the PL-approximability.

Another consequence of having a biological function is that the spectrum of a genome has a non-uniform distribution. To test the effect of this, we process the *k*-mers of chr1 with a hash function and use the hashed value as the basis for the rank function. This has the effect of making the distribution uniform. The resulting *α*^∗^ = 1.43 shows that the non-uniform distribution of the *k*-mers in a spectrum substantially decreases the PL-approximability.

Finally, we combined the two effects by removing repeats and then hashing the *k*-mers, getting an *α*^∗^ = 1.87, the same as a random multiset. This indicates that the two factors have an interplay which drives down the PL-approximability a lot when combined.

We repeated the same experiments on the Y chromosome (Figure 8). The effects were similar, though the effect of removing repeats was more pronounced (*α*^∗^ rose from 0.96 to 1.09). This was expected, as the Y is known to be more repetitive than chromosome 1 [28].

### 5.5 Stability of CaPLa as a measure

#### Uniqueness of pinch point

We conducted brute-force tests to investigate whether in our data the pinch point is unique (i.e. CaPLa is well defined), and to verify whether the unimodality assumption required by our algorithm in Theorem 10 holds in practice. Specifically, for each genome in the RefSeq dataset, we sampled the twisted ribbon width *W* (*α*) at uniformly spaced *α*-values within the interval [*α*_*L*_, *α*_*H*_] containing a pinch point (Theorem 9), using a step size of 10^−6^. We then checked for the presence of multiple minima within a numerical tolerance of *τ* = 1.49×10^−8^, and verified unimodality by looking for a decreasing and increasing trend before and after the minimum, respectively.

We found that the pinch point was unique in all of the 513 RefSeq genomes. Furthermore, the unimodality assumption was not met in only 11 datasets. Even in those 11 cases, the algorithm of Theorem 10 avoided local minima and matched the brute-force pinch point in all but one instance, where the actual pinch point was nonetheless close (within 3.23×10^−2^). In summary, up to the tested step size, CaPLa was defined for all genomes and our efficient algorithm returned the correct value in all but one case.

#### Genome length

A desirable property of a measure of PL-approximability is that it is not causally dependent on genome length and is only an intrinsic property of a genome’s rank curve. To verify this, we compare the lengths of our RefSeq genomes with their *α*^∗^, 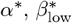, and 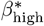 values, respectively, as well as to the *ε*-mapped CaPLa, for *ε* ∈ {2, 4, 8, …, 1024}. The Pearson correlation coefficient (*R*^2^) is less than 0.02 for all comparisons, providing evidence that the relationship between CaPLa and genome length is negligible.

#### *k*-mer size

The structure of the rank curve is unavoidably dependent on *k*, because for small *k*, the number of repeats and the density of *S* with respect to the universe gets high. Ideally, when *k* is above the threshold where spurious repeats are rare, CaPLa should stabilize. To validate this, we looked at the behavior of CaPLa with respect to *k* for the RefSeq dataset and for bigger mammalian genomes. Figure 9 shows six genomes, which we manually selected to capture the phylogenetic diversity of the data. We measure the averaged *ε*-mapped CaPLa for *ε* = 64. The plots show that indeed CaPLa plateaus at some point between *k* = 16 and *k* = 21, depending on the genome. The difference between the averaged *ε*-mapped CaPLa values at *k* = 26 and *k* = 31 was less than 0.01 for all six genomes, both for *ε* = 64 (Figure 9) and *ε* = 32 (data not shown). Similarly, between *k* = 21 and *k* = 26, the differences in *α*^∗^ values were less than 0.4% and the difference in the 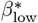 and 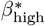 values were less than 4%. These results provide strong evidence that the CaPLa is relatively stable for large enough values of *k*, and, in particular, for values of *k* ≥ 21.

**Figure 9.**
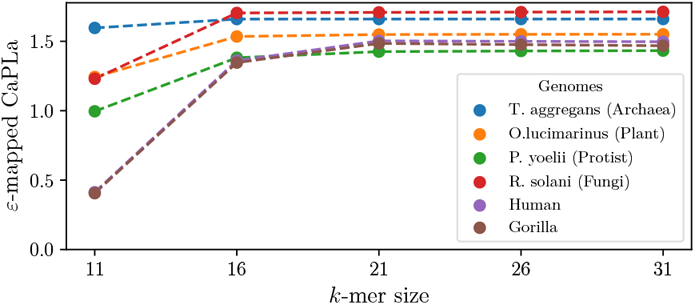
Effect of *k*-mer size on CaPLa, for *ε* = 64, using averaged *ε*-mapped CaPLa

#### Accuracy of prediction

We compare the space range predicted by plugging in CaPLa into Theorem 11 with the true space taken by the PLA-index (Figure 10). In the majority of cases, the true space falls within the predicted range, as expected. However, for most cases of *ε* = 1024 and one case of *ε* = 16, the true space is higher than the predicted range. This is likely due to one of two reasons: 1) the PLA-index uses an implementation of Elias-Fano encoding that has a constant overhead per character, which is not the theoretically optimal data structure used for theoretical analysis, and 2) the results of Theorem 11 hold only asymptotically and do not account for constant factors that contribute to the space, especially for higher *ε* where the total space is small.

**Figure 10.**
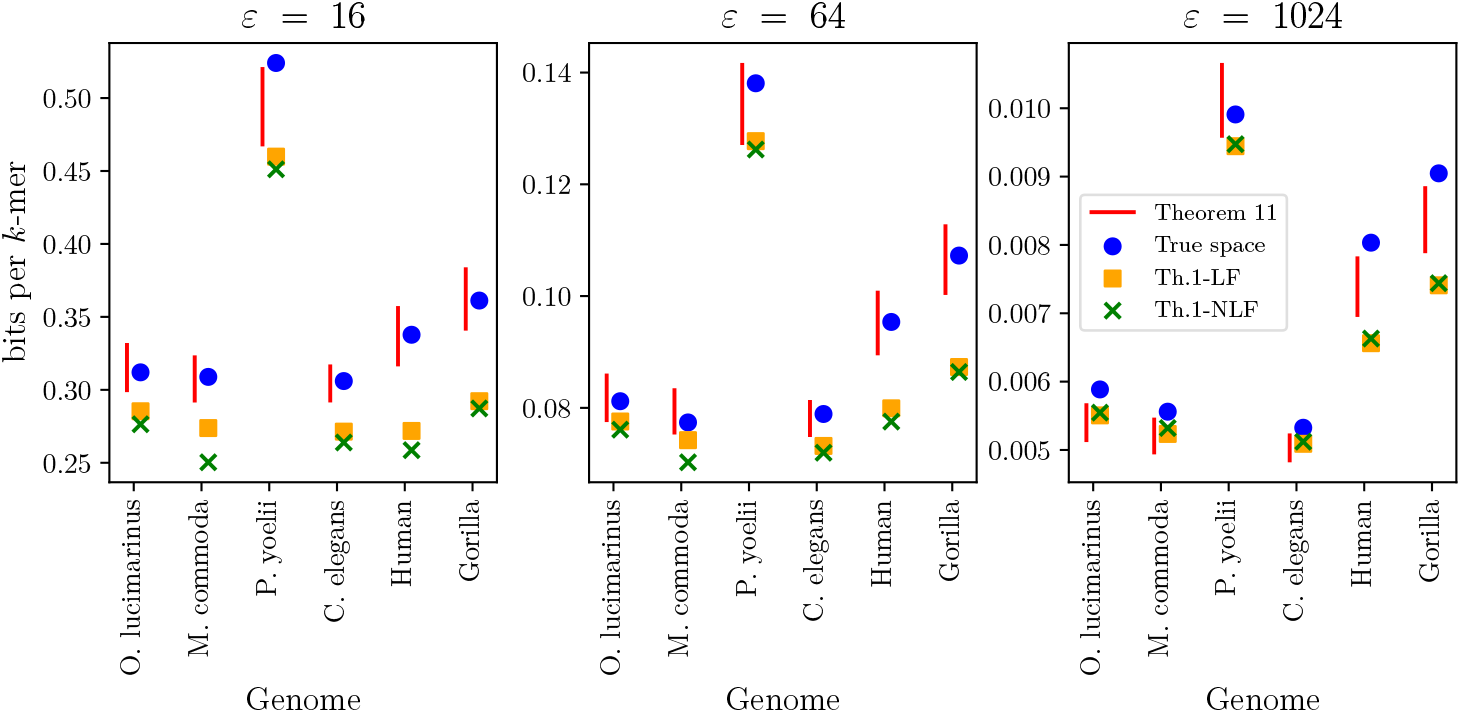
The true space of the PLA-index compared with various forms of predictions, for *ε* ∈ {16, 64, 1024}. The true space is measured using a version of the PLA-index implementation which encodes the X-array using Elias-Fano, which is the version that is theoretically analyzed in [1] and in Theorem 11. The predicted range (red) corresponds to plugging in the CaPLa into Theorem 11, using *c* = 7. The Th.1-LF and Th.1-NLF correspond to fitting a power-law model with a linear and non-linear fit, respectively, and plugging in the fitted parameters into Theorem 1 in [1] (with *c* = 4). The fitted and non-fitted models are described in the caption of Figure 2.

We also compare the true space with the space predicted by fitting a single power-law model to the tabulated PLA-sizes (as we previously discussed in Figure 2) and plugging the fitted parameters into Theorem 1 from [1], with *c* = 4. As Figure 10 shows, the fit-based predictions are often significantly under-predicting the true space, especially at *ε* = 16.

## 6 Conclusion

In this paper, we introduced the notion of PL-approximability of a multiset and a corresponding measure called CaPLa (Section 2). Our theoretical results show that CaPLa is well-defined and can be computed efficiently, under reasonable assumptions which we demonstrate hold in almost all tested data (Section 3). Our experimental results show that CaPLa varies substantially not only across genomes and kingdoms, but also within individual genomes, and this variability can lead to differences of up to 50% in the space required by a PLA-based data structure (Sections 5.2 and 5.3). Within our CaPLa framework, the key factors distinguishing a genome spectrum from a random *k*-mer multiset are the presence of repeats and the non-uniform distribution of *k*-mers. The combined effect of these two factors substantially reduces PL-approximability (Section 5.4). CaPLa has several desirable properties, namely, it is well-defined and efficient to compute across our experimented genomes, it is not trivially correlated with genome length thus indicating it captures the intrinsic properties of the genome, and it is robust with respect to *k*-mer size when *k* ≥ 21 (Section 5.5).

Our result has a concrete application to parameter selection in PLA-based indexes such as the PLA-index. When a target space is specified, an approximate estimate of the dataset’s CaPLa can be used with Theorem 11 to compute a suitable value for the tuning parameter *ε*. This is particularly advantageous when the CaPLa is known for a similar dataset (e.g., from another individual of the same species), as it avoids the need to recompute the full set of (*ε, b*(*ε*)) pairs. In the absence of prior CaPLa information, future work could also explore efficient estimation of CaPLa over a small sample of the dataset itself (e.g., a contiguous region of the suffix array).

Our paper presents several future directions. From an application perspective, it will be interesting to connect the variability of PL-approximability to biological properties. Our preliminary results show that PL-approximability is higher in fungi and lower in repeat-rich chromosomes like the Y, but this raises more questions than it answers. From a theory perspective, it would be interesting to find the conditions under which the pinch point is unique. In our experiments, we could not find any instances of non-unique pinch points, suggesting that a theoretical proof may be possible. Finally, Theorem 9 requires that 1∈ ε in order to guarantee the existence of a pinch point. This condition is necessary, as in fact one can prove that a pinch point does not exist if 1 ∉ε. This raises an interesting open question if there is some alternative notion of CaPLa that can be used when 1 ∉ε.

## Acknowledgements

We thank anonymous reviewers for helping improve various versions of this paper.

---

**Figure A.1.**
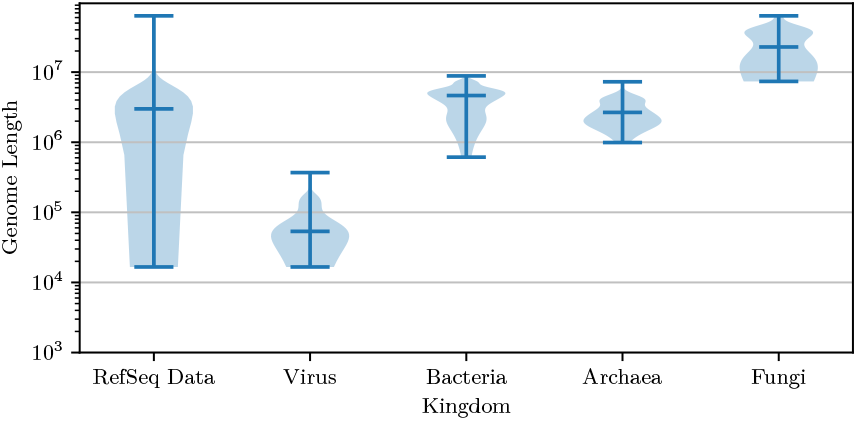
Genome length distribution in RefSeq, shown for the whole dataset and separately for each of the four constituent kingdoms. The three horizontal lines indicate the minimum, median and maximum values.

**Table A.1.** Length statistics for six additional genomes not in our RefSeq dataset for *k* = 21.

| Genome | $n$ (mil) | $N$ (mil) |
| --- | --- | --- |
| *C. elegans* | 93 | 100 |
| *P. yoelii* | 20 | 23 |
| *M. commoda* | 21 | 21 |
| *O. lucimarinus* | 13 | 13 |
| Human | 2,333 | 3,117 |
| Gorilla | 2,346 | 3,595 |

## A Appendix

This Appendix contains figures and proofs omitted from the main text due to space constraints.

### ▸ Lemma 8

*If* 1 ∈ ε *and* |ε|*>* 1, *then:*

i. *H and L are continuous functions*.
ii. *For any α* ≥ 0, *it holds* 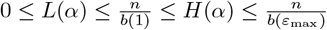, *where ε*_max_ = max E.
iii. *There exists a value α*_*L*_ *such that* 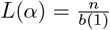, *and L*(*α*_*L*_ + *δ*) *decreases as δ>* 0 *increases. We call α*_*L*_ *a flattening point of S*.
iv. *There exists a value α*_*H*_ *such that* 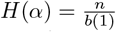, *and H*(*α*_*H*_ − *δ*) *increases as δ>* 0 *increases. We also call α*_*H*_ *a flattening point of S*.
v. *The flattening points satisfy α*_*L*_ ≤ *α*_*H*_.

**Proof**. Let 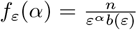, thus *L*(*α*) = min_*ε*∈ ε_ *f*_*ε*_(*α*) and *H*(*α*) = max_*ε*∈ ε_ *f*_*ε*_(*α*). Observe that *f*_*ε*_(*α*) is a continuous function of *α* for any fixed *ε*. The min or max of a set of continuous functions is itself continuous, which proves i.

Note also that differentiating *f* _*ε*_(*α*) with respect to *α* yields 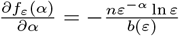. If *ε* = 1,the derivative is equal to 0 since ln *ε* = ln 1 = 0. For all other *ε >* 1, the derivative is strictly negative, since *b*(*ε*) and the terms in the numerator are positive. Hence *f*_*ε*_(*α*) is a decreasing function of *α* if *ε* ≠ 1, and constant when *ε* = 1.

To prove iii, observe first that 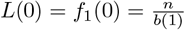. Moreover, 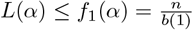, since by definition *L*(*α*) is the minimum 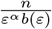 over ε. By assumption, | ε | *>* 1, so there exists at least a value *ε*^′^ ≠ 1 in ε. From the discussion above, *f*_*ε*_′ (*α*) is a decreasing function and is such that *f*_*ε*_′ (*α*) → 0 as *α* → ∞. Thus, by the intermediate value theorem, there exists a value *α*_*L*_ at which the function 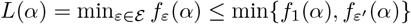 transitions from being the constant 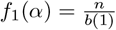 to decreasing to 0 as *α* → ∞.

To prove iv, observe first that 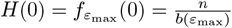. Moreover, 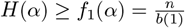,since by definition *H*(*α*) is the maximum 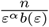 over ε. Similarly to the previous point,there exists an *ε*^′^ ∈ ε \ {1} and a value *α*_*H*_ at which the function *H*(*α*) = max_*ε*∈ ε_ *f*_*ε*_(*α*) ≥max{*f*_1_(*α*), *f*_*ε*_′ (*α*)} transitions from being equal to the function *f*_*ε*_′ (*α*), which decreases to 0 as *α* → ∞, to being the constant 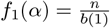

To prove v, suppose for the sake of contradiction that *α*_*H*_ *< α*_*L*_. For all *α* ∈ [*α*_*H*_, *α*_*L*_], it holds 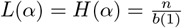 due to iii and iv. From Lemma 6, we also have *L*(*α*) ≤ *f*_*ε*_(*α*) ≤*H*(*α*) for any *ε* ∈ ε, which, combined with the previous equality, implies 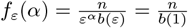for any *ε* ∈ ε.

Now, consider any *ε* ∈\{1}and pick two distinct values *α*_1_, *α*_2_ ∈ [*α*_*H*_, *α*_*L*_]. For these values, the last equality implies

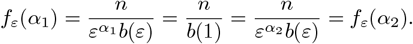

Simplifying, this means *ε*^*α*1;^ = *ε*^*α*2;^, which is impossible since *α*_1 ≠_ *α*_2_ and *ε ≠* 1. Therefore, we conclude that *α*_*L*_ ≤ *α*_*H*_.

Finally, observe that ii is a consequence of iii and iv.

### ▸ Theorem 9

*If* 1∈ε *and* |ε | *>* 1, *then a pinch point α*^∗^ *of S exists and is guaranteed to lie in the interval* [*α*_*L*_, *α*_*H*_], *where α*_*L*_ *and α*_*H*_ *are the flattening points of S*.

**Proof** Consider the interval [*α*_*L*_, *α*_*H*_], which is well-defined due to Lemma 8v. We show that the twisted ribbon width *W* (*α*) = *H*(*α*)−*L*(*α*) can only increase outside of this interval, from which we conclude that *α*^∗^ must occur within this interval. The existence of *α*^∗^ within this interval is guaranteed by the extreme value theorem.

Let us first consider values of *α* ≥ *α*_*H*_ by setting *α* = *α*_*H*_ + *δ*for some *δ*≥ 0. By Lemma 8iv, 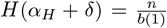 for any *δ*≥ 0. By Lemma 8iii, *L*(*α*_*H*_ + *δ*) decreases as *δ*increases. Therefore, 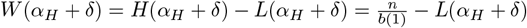 increases as *δ*increases.

Let us now consider values of *α* ≤ *α*_*L*_ by setting *α* = *α*_*L*_ − *δ*for some *δ*≥ 0. By Lemma 8iii, 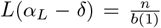 for any *δ*≥ 0. By Lemma 8iv, *H*(*α*_*L*_ − *δ*) increases as *δ*increases. Therefore 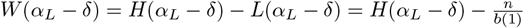 iincreases as *δ*increases.

▸ **Theorem 11***Let S be a multiset of k-mers, with n distinct elements, N total elements. Suppose the PL-approximability of S is* (*α, β*_low_, *β*_high_) *power-law bounded over* ε. *Let B*_*ε*_ *be the number of bits used by the PLA-index on S with tuning parameter ε. Then, for all ε, we have that*

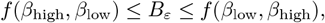

*Where*

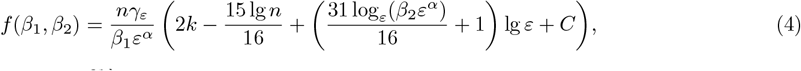

*and* 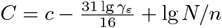 *and, for large enough* 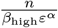, *c is a number between 5 and 7*.

**Proof**.Theorem 1 in [1] says that the space used by PLA-index with its default parameter *l* = 1*/*16 is

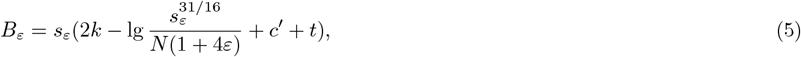

where *c*^′^ is a constant between 3 and 4 and *t* denotes a positive term that goes to 0 as *s*_*ε*_ → ∞, which implies that *t* → 0 as *b*(*ε*) → ∞.

Next, we can use the definition of power-law bounded PL-approximability to sandwich lg *b*(*ε*). For convenience, let *ρ*_low_ = *α* + log_*ε*_ *β*_low_ and *ρ*_high_ = *α* + log_*ε*_ *β*_high_.

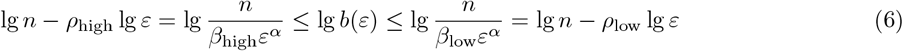

Note that this implies that *t* → 0 as 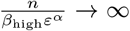. We will need the following bounds on lg(4*ε* + 1).

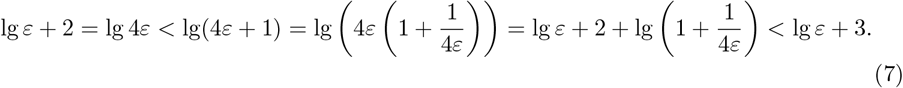

The first and last inequalities use the fact that *ε* ≥ 1. Therefore, for large enough 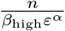, lg(4*ε* + 1) + *c*^′^ + *t* lies between lg *ε* + 5 and lg *ε* + 7.

Plugging these facts into *B*_*ε*_, we get

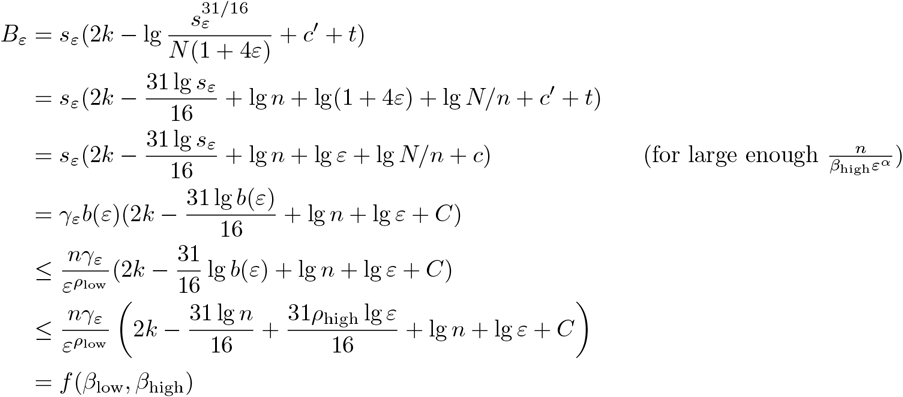

The lower bound on *B*_*ε*_ is derived in a symmetric way, with the roles of *ρ*_low_ and *ρ*_high_ reversed as in Eq. 6.

## Footnotes

1 Our tool for computing CaPLa is available at https://github.com/medvedevgroup/CaPLa.

